# Persistence of cooperation in diffusive public goods games

**DOI:** 10.1101/352492

**Authors:** Philip Gerlee, Philipp M. Altrock

## Abstract

Diffusive public goods (PG) games are difficult to analyze due to population assortment affecting growth rates of cooperators (producers) and free-riders. We study these growth rates using spectral decomposition of cellular densities, and derive a finite cell-size correction of the growth rate advantage, which exactly describes the dynamics of a randomly assorted population, and approximates the dynamics under limited dispersal. The resulting effective benefit to cost ratio relates the physical parameters of PG dynamics to the persistence of cooperation, and our findings provide a powerful tool for the analysis of diffusive PG games, explaining commonly observed patterns of cooperation.

## INTRODUCTION

Many organism are known to produce costly goods (e.g. molecules) that are released into the surrounding environment. These goods provide a benefit not only to producers themselves, but also to others. Examples of this behavior range from microorganisms such as bacteria that release digestive enzymes [1] and siderophores that aid in the uptake of iron [2], to plants that release compounds that reduce herbivory [3], and beetles that produce pheromones that increase reproductive success [4]. Since public good production comes at a cost, the resulting population dynamics are an evolutionary game.

Public goods (PG) games are also important in cancer biology. In evolving tumors growth factors (a type of hormone) typically produced by tumor associated cells (e.g. fibroblasts), have the ability to increase the rate of tumor growth [5, 6]. Recent findings have shown that tumor cells can acquire the capability of producing growth factors which benefit themselves and neighboring cells [7]. This includes PDGF-production in brain cancer cells [8], Wnt-production in breast cancer [9] and testosterone-based interactions in prostate cancer cells [10]. Tumor cells typically harbor mutations that may lead to the production of a public good in form of a secreted factor. Production often comes at a fitness cost. Hence one can ask how the production of a costly public good can persist [11]. To answer this question one needs to examine the relation between benefits and costs incurred by PG production. Only if the average benefit outweighs the cost, can producers potentially spread in the population.

There have been several attempts to understand the problem of PG producer persistence in relation to the individual cost. In an important set of experiments, Archetti *et al.* showed that, depending on external conditions, either production or free-riding is favored [11]. For intermediate background serum concentrations the two cell types could coexist, but either type dominated in the case of low and high serum concentrations. This phenomenon can be explained using evolutionary game theory. A drawback with many game theoretic approaches is an assumed time-scale separation between interaction and reproduction [12]. Fitness values are calculated by averaging over configurations of individuals [13, 14], which is necessary as the PG is a multi-player game, but could be problematic since the total number of feasible configurations might be limited and biased due to spatial correlations, especially for limited cell movement and dispersal. The actual size of the local group (or a distribution thereof) might be impossible to determine from experiment [11, 15].

General condition for persistence of cooperation for different spatial arrangements of cells or microbes in 2D were obtained by Allen *et al*. [16]. A potential drawback of this approach is that PG diffusion was assumed to occur on a network; PG is redistributed only to nearest neighbors on the network or lattice. This implies that the diffusion parameter does not necessarily have a direct physical meaning, but rather quantifies the amount of molecules that are transferred along edges of the population graph. Similarly, an exact condition for when selection favors public good producers in a network-based diffusible PG game was derived by Driscoll and Pepper [17], who sought to capture the evolutionary game’s dependence on the diffusion parameter, which determines the extent to which the PG is retained rather than distributed to neighboring cells. But also in this case it is difficult to relate the theoretical predictions to actual experiments. In *in vitro* and *in vivo* experimental systems, PG diffuses further than nearest neighbors and influences more distant individuals. This long-range phenomenon would have to be captured by an alternative graph structure that cannot be determined ad hoc, and would require weighted links.

Directly modeling of diffusion based dynamics of PG within a spatially distributed population of cells that interact mechanically represents a more realistic approach [18]. Using such a model it was shown that the spatial distribution of genetic lineages and the population dynamics of producers and free-riders is linked to cell growth rate, nutrient availability, and nutrient diffusivity [19]. In a computational study of biofilm production by Xavier and Foster [20], it was shown that there is a strong evolutionary advantage to extracellular polymer production. Secretion of polymers is altruistic to cells above a focal cell since it pushes later generations in their lineage up and out into better nutrient conditions, but it harms others because polymer production tends to suffocate neighboring non-polymer producers.

We here seek to understand how key parameters of the public good, such as its diffusion coefficient, its production rate, and the cost, affect co-evolution and growth of producers and free-riders. We consider a system of intermediate complexity, between the simple network-based approach and the analytically intractable biophysical systems. We do not a priori assume that the public good induces frequency-dependent selection, or an implicit evolutionary game based on individual payoffs. Instead, we take a bottom-up approach and explicitly model production, diffusion and decay of the public good, as well as the spatial distribution of producers and free-riders, in order to describe how these distributions change over time. We establish analytical predictions that quantify under which conditions cooperation in the form of PG production can prevail.

## RESULTS

### Individual-based model

We start our investigation with an IB-model in which the cells reside on a two-dimensional square lattice (see fig. 1 (a) and Supplement for details). The linear size of the domain is *L* = 1 cm and it contains *N × N* sites separated by a distance *L/N,* which will be a key quantity in our analysis. For cancer cells a meaningful value is *N* = 200, which yields a cell radius of 25 *µ*m [21]. The PG concentration evolves according to

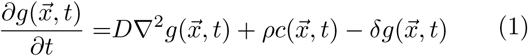

where 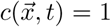 at all sites that contain producers and zero otherwise. The public good is subject to no-flux boundary conditions representing a closed experimental system. A free-rider cell located at site 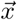 divides at a rate 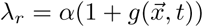, whereas a producer cell pays a fixed cost *κ* and hence divides at rate *λ*_*c*_ = *λ*_*r*_ − *κ*. Upon cell division the daughter cell is placed uniformly at random in a neighboring site. If the site is occupied cell division fails. Cells of both types are assumed to die at a constant rate *µ*, and also move into empty neighboring sites at rate *V*. These processes give rise to a complex dynamical system, for which fig. 1 (b) and (c) show snapshots of the producer and free-rider cell densities and the resulting distribution of public good, respectively. All parameters and their values and units are given in Table I.

**TABLE 1.**
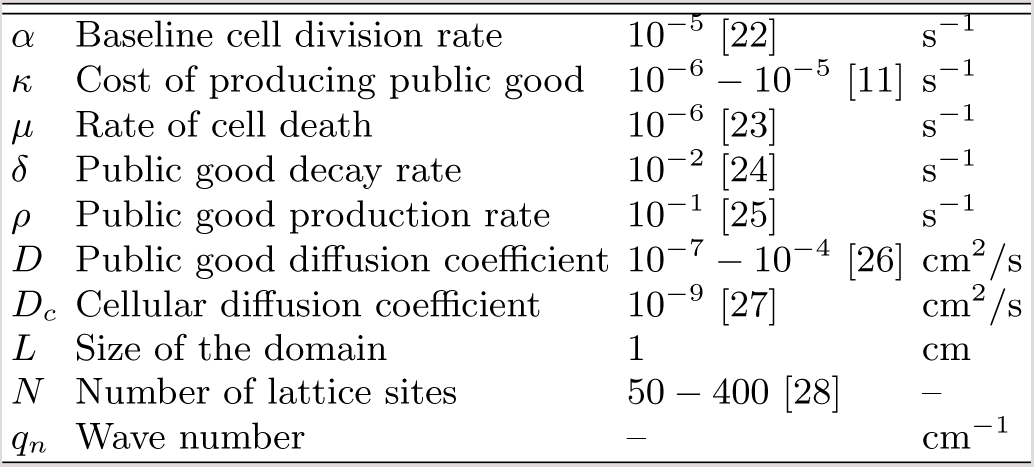
Model parameters

|  |  |  |  |
| --- | --- | --- | --- |
| $\alpha$ | Baseline cell division rate | $10^{-5}$ [22] | $s^{-1}$ |
| $\kappa$ | Cost of producing public good | $10^{-6} - 10^{-5}$ [11] | $s^{-1}$ |
| $\mu$ | Rate of cell death | $10^{-6}$ [23] | $s^{-1}$ |
| $\delta$ | Public good decay rate | $10^{-2}$ [24] | $s^{-1}$ |
| $\rho$ | Public good production rate | $10^{-1}$ [25] | $s^{-1}$ |
| $D$ | Public good diffusion coefficient | $10^{-7} - 10^{-4}$ [26] | $cm^2/s$ |
| $D_c$ | Cellular diffusion coefficient | $10^{-9}$ [27] | $cm^2/s$ |
| $L$ | Size of the domain | 1 | $cm$ |
| $N$ | Number of lattice sites | 50 – 400 [28] | – |
| $q_n$ | Wave number | – | $cm^{-1}$ |

**FIG. 1.**
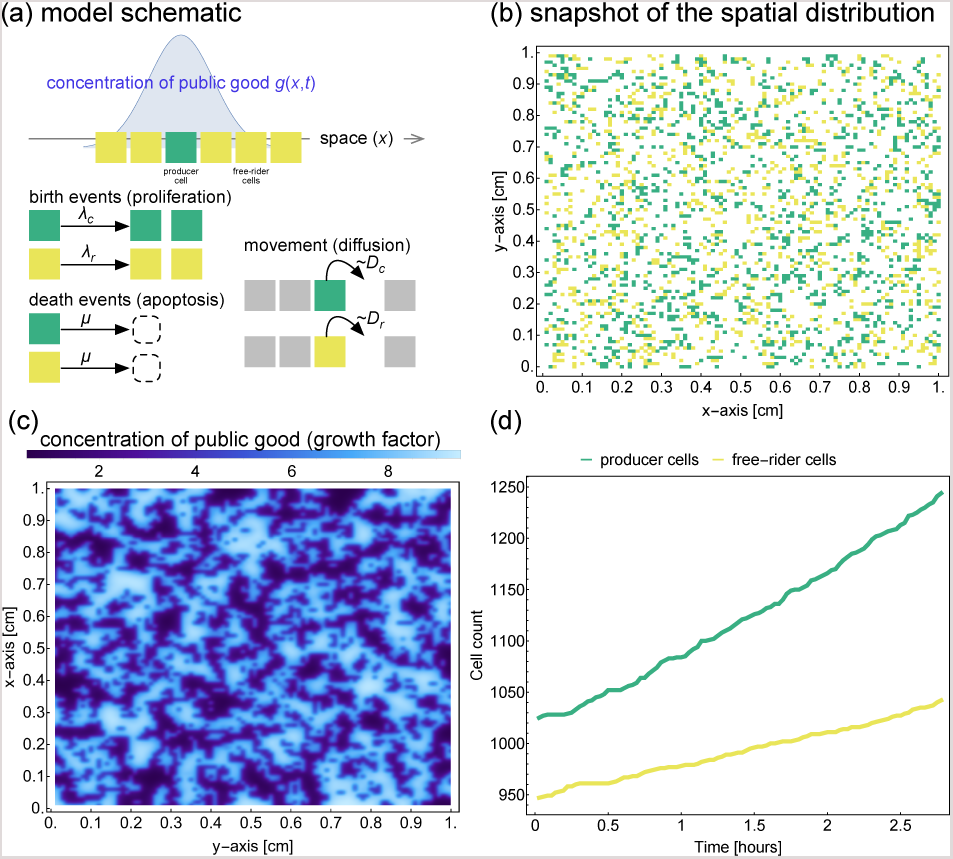
(a) Schematic of the individual-based model with a diffusible public good public good *g*(*x, t*). Cells are located on a lattice and can be of two types: producers cells that produce the diffusible public good, and free-riders that do not produce but still receive benefit in the form of increased rate of cell division. The public good is produced at rate *ρ*, diffuses with diffusion constant *D* and decays at a constant rate *δ*. A schematic of the concentration in space is given as the solid curve, i.e. the PG concentration spikes near producer cells. In a typical simulation of the model the cells divide and place their offspring at neighboring lattice sites. This changes the spatial distribution public goods concentration (shown in c), which in turn affects the growth curves of the two cell types (shown in d).

For a given initial distribution of cells this models makes it possible to simulate the population dynamics and for a given parameter configuration determine if producers or free-riders dominate (see fig. 1 (d)). However, this method is computationally expensive and we have therefore aimed for an analytical treatment of the problem.

### Continuum limit approach

We considered the following continuum approximation to describe the dynamics of the cellular densities in space and time, in the limit of negligible cell size (see Supple-mental Methods):

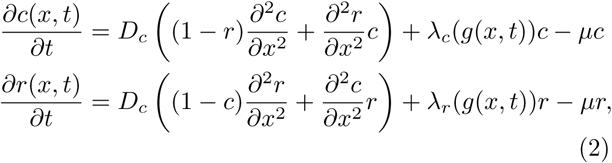

where *c*(*x, t*) and *r*(*x, t*) denote the normalized density of producer cells and free-riders respectively. Here we consider a 1-dimensional system since it simplifies calculations, but still yields predictions applicable to the 2D-model. The public goods concentration *g*(*x, t*) varies in both space and time and its dynamics are driven by production, diffusion and decay. A separation in time-scale between PG dynamics and cell division implies that we could assume the following stationarity condition for the PG concentration:

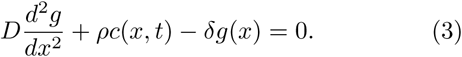

If such a time-scale separation is appropriate, the public good concentration will explicitly be driven by the spatial distribution of PG producer cells.

### Logistic growth model

In order to quantify how producer cells fare over time, we disregard the spatial structure and instead analyze changes in the total number of producer and free-riders cell, which are given by

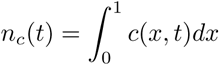

And

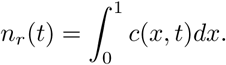

Next we seek to quantify how these quantities change over time (see Figure 1 (d)). Under the assumption of long-range dispersal of daughter cells the dynamics of the population densities can be approximated by a system of coupled logistic equations [29]:

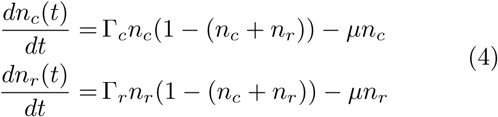

where Γ_*c*_ and Γ_*r*_ are the average per capita growth rates, which depend on the spatial distribution of PG, and remain to be calculated. In the case of short range dispersal as in our IB-model a higher order correction term, which accounts for spatial correlations, can be derived [29]. However, this correction only has a minor impact on the long-term dynamics, and the logistic system provides an accurate description of our system.

We next turn to a derivation of the average growth rates Γ_*c*_ and Γ_*r*_. This derivation contains a number of intermediate steps: First, we show that by decomposing the spatial density of cells into Fourier coefficients it is possible to calculate how these coefficients impact the instantaneous growth rates. Cell birth and death will cause the Fourier coefficient to evolve in time, and although it is possible to write down equations for these dynamics we have taken a different approach here. Second, we describe the discrete and stochastic nature of the cell densities in the IB-model (see Figure 1 (b)) as a space-discrete stochastic processes. This allows us to describe a benefit to self experienced by producer cells, which cannot be captured by the continuum approach. Third, we show that the relation between Fourier coefficients and growth rates can be extended to the stochastic model, and explicitly derive the expected growth rate for a population undergoing long-range dispersal. The key to this is the realization that if the model with long-range dispersal is initialized without any spatial structure then as time passes the densities will change, but no spatial correlations will appear. This implies that the expected values of the Fourier coefficients of the cellular densities will only depend on the densities, and consequently the expected growth rates Γ_*c*_ and Γ_*r*_ will also exclusively depend on the densities, thus eliminating the dependence on the PG distribution, and turning (4) into a closed system of equations.

### From Fourier coefficients to growth rates

How can we approximate the growth rates of producers and free-riders in order to evaluate the success of cooperators in the system? In the Supplementary Methods we prove a theorem that relates the parameters of the model to the growth rates in an exact way under specific geometric assumptions regarding the spatial distributions of producers and free-riders. As a result, we find that if the densities *c*(*x, T)* and *r*(*x, T)* are both square integrable, then the growth rates of producer and free-rider cells at time *t* = *T* are given by

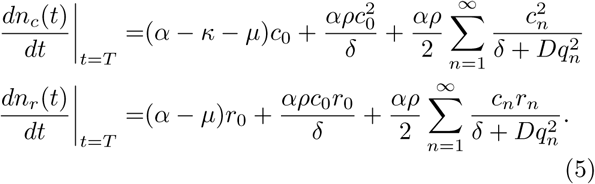

where *q*_*n*_ = 2*πn/L*, and *c*_*n*_ and *r*_*n*_ are the coefficients of the cosine Fourier series of *c*(*x, T)* and *r*(*x, T)* respectively. This result states that the growth rate of producer cells is a linear function of the cost of PG production, and that both growth rates are linear in the production rate, and establishes a reciprocal relationship to both the PG diffusion constant and the wave number, which describes the scale of spatial oscillation. A key assumption underlying the proof of this relationship is that the cellular densities can be represented as Fourier series on the unit interval.

We next calculated the expected growth rates for arbitrary stochastic initial cellular distributions. Stochastic, uncorrelated cellular distributions are realistic if the cell types were subject to a well-mixed solution before plating occurred. Subsequent cell division and long-range dispersal maintains the uncorrelated spatial structure. We model this by prescribing the initial condition on the lattice according to Bernoulli processes, where occupation probabilities between sites are uncorrelated, and for each lattice site *p*_*c*_ is the probability to find it occupied by a producer cell, *p*_*r*_ is the probability to find it occupied by a free-rider cell and it is empty with probability 1*−p*_*c*_*− p*_*r*_ (See Supplementary Methods for details).

In the continuum limit such a well-mixed initial condition corresponds to initial densities that are constant in space, *c*(*x,* 0) = *c*_0_ and *r*(*x,* 0) = *r*_0_, for some positive constants *c*_0_ and *r*_0_. Consequentially, all other Fourier coefficients would be zero. In this case, the result (5) tells us that the growth rates should be independent of the diffusion coefficient *D*. However, this parameter independence is not what we observe in the stochastic individual based simulations with initially homogenous distributions of cells in space. Instead, the *expected* growth rates, after stochastic plating, depend on the PG diffusion coefficient *D* (data not shown). This behavior is captured if we model the initial cell densities as two dependent Bernoulli processes with parameters *p*_*c*_ and *p*_*r*_. The expected growth rates can be calculated as (see Sup-plement for details)

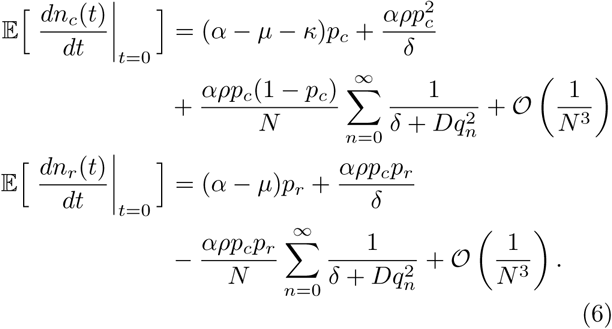

where *N* is the total number of lattice sites in the domain, which is inversely related to the cell size. The infinite sum can be written as

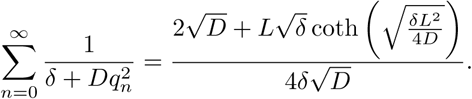

Here, the term 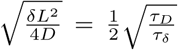, where *τ*_*D*_ = *L*^2^*/D* is the diffusion time scale and *τ*_*δ*_ = 1*/δ* is the time scale of decay. For realistic parameter regimes, we expect that *τ*_*D*_ *≫ τ* _*δ*_ which implies that coth 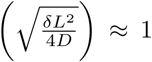. This means that

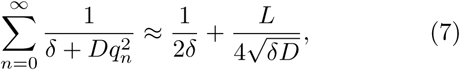

which simplifies further calculations.

### Predicting evolutionary outcomes

With the theory developed so far we can now obtain first order finite-size corrections of the average growth rates Γ_*c*_ and Γ_*r*_. In the limit of cell size tending to zero (*N → ∞*) the expected growth rates given in (6) are equal to the continuum result given in (5), with *p*_*c*_ = *c*_0_, *p*_*r*_ = *r*_0_ and *c*_*n*_ = *r*_*n*_ = 0 for *n ≥* 1, as argued above. The expected growth rates (6) give the average rate of cell division and we can therefore equate them with the growth rates (4),

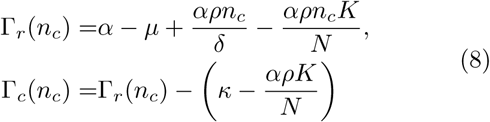

with the constant

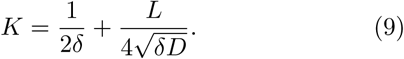

These two growth rates only depend on the density of producers cells, and the difference is 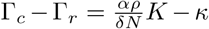,which can be interpreted as the self-benefit to producer cells that scales with the cell size (1*/N)*, minus the cost of production *κ*. This results is only exact (to first order in 1*/N)* for a system with long-range dispersal, but agrees well with the IB-model simulations under local dispersal.

An example of a numerical solution of the logistic system (4) with growth rates calculated according to (8) is shown in Figure 2 (a,b), where the dashed lines correspond to the dynamics of the IB-model and the solid lines correspond to the logistic system. The discrepancy is largest early on when local dispersal limits the population growth. However, when the system has reached its carrying capacity the dynamics are well predicted by the logistic system. A systematic exploration of the agreement between the logistic system (analytical) and the IB-model (simulations) is shown in Figure 2 (c); the theoretical prediction is within 5 % of the IB-model as the diffusion coefficient is varied across 3 orders of magni-tude.

**FIG. 2.**
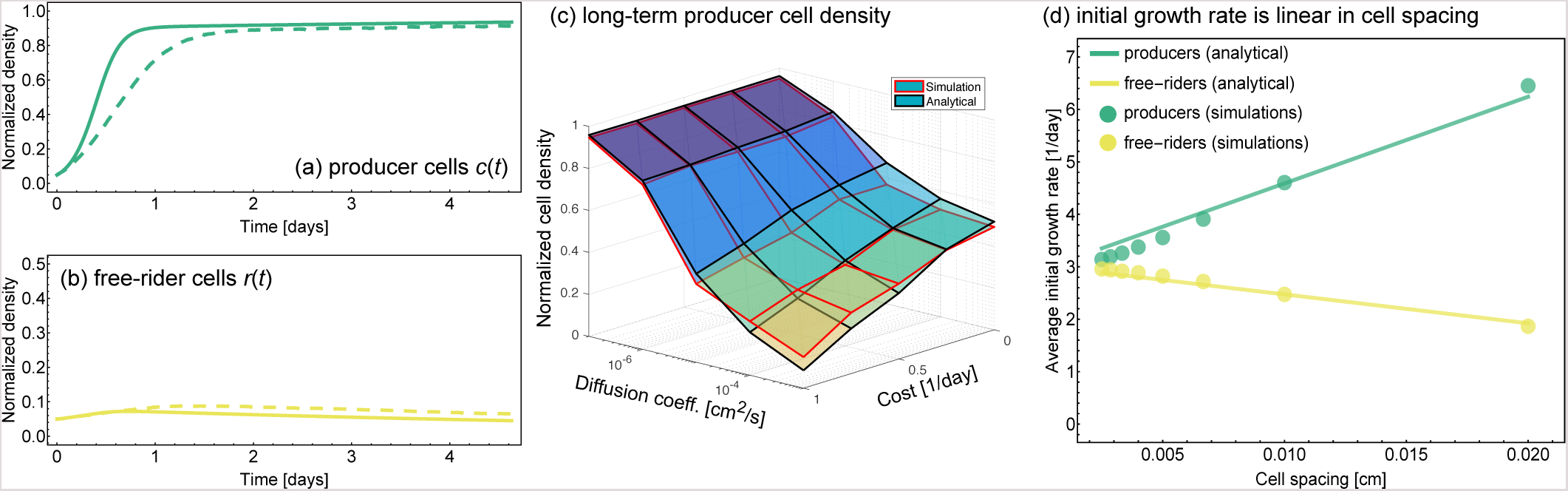
The dynamics of the IB-model with local dispersal. **(a,b)** Using our analysis of stochastic and uncorrelated spatial structure it is possible to predict the long-term dynamics of the individual-based model using the set of coupled logistic equations (4). The solid lines correspond to the solution of Eq. (4) and the dashed lines are obtained from a single run of the IB model. The density of public goods producers after 10 days as a function of the public good diffusion coefficient and the cost of production. The initial densities of producers and free-riders were set to *p*_*c*_ = *p*_*r*_ = 0.05, i.e. each point in space had an equal chance to be occupied with a producer or free-rider cell. The surface with red edges corresponds to results obtained by averaging 10 simulations of our individual-based model, and the surface with black edges was obtained by solving the system (4) numerically. **(d)** The average initial growth rate as a function of the spacing between the cells, when cells are seeded at initial density *n*_*c*_(0) = *n*_*r*_ (0) = 0.05. The results were obtained by averaging over 10 simulation for each value of the cell spacing. As predicted, the growth rates depend linearly on the cell spacing 1*/N,* and closely follow the theoretical prediction (8) (solid lines). All parameter values are given in Table I, with the PG diffusion constant *D* = 10^*−*6^ cm^2^/s (unless indicated otherwise) and the cost *κ* = 10^*−*6^ s^*−*1^.

Analysis of the logistic system reveals that long-term co-existence cannot be observed (see Supplemental Methods for details). The perceived co-existence, shown in fig. 2 (a,b), is thus transient; in the long run free-riders will go extinct for this parameter configuration. The long-term evolutionary outcome is determined by stability of the monomorphic fixed points, which are controlled by the sign of the effective cost of PG production

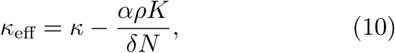

where *K* is given by (9). If *κ*_eff_ *>* 0, producers will out-compete free-riders, otherwise the free-riders will dominate.

The theoretical prediction (8) states that the average growth rates scale as 1*/N,* which is proportional to cell spacing. In simulations, we measured the average growth rate of producers and free-riders when seeded randomly with a fixed density and varied the cell spacing. The comparison is shown in figure 2 (d), which shows that our theoretical prediction closely matches the individual-based simulations, and confirms that the average growth rates depend linearly on the spacing (1*/N)*. The discrepancy observed for larger spacing (small *N)* is due to the fact that the continuum model poorly describes the system for small *N*.

## DISCUSSION

We have investigated the dynamics of a population of organisms made up of two distinct types, producers that release a diffusible public good (PG) into the environment, and free-riders. Starting with an individual-based model of the system we derived a set of coupled PDEs that describe how the densities change in space and time. We showed that the growth rates of the two types can be expressed in terms of Fourier coefficients, highlighting that spatial oscillations impact the dynamics in a wavelength dependent manner. This approach also made it possible to obtain results for the case of a randomly distributed population, showing that producer cells obtain a benefit that depends on the cell size, effectively describing the amount of PG retained by the cells themselves. We also showed that the long-term dynamics are determined by the sign of an *effective* cost of cooperation, for which we considered a well-mixed approximation of the spatial system. If the effective cost is positive producers will outcompete free-riders. This result is exact for long-range dispersal, but also provides a good approximation for local dispersal.

Existing theoretical results are derived from models that fail to capture the actual reaction-diffusion dynamics of the public good in the spatial domain [16, 17], whereas more realistic models are typically too complex to analyze mathematically [18]. Our results represent a novel contribution to the study of spatial public good games since we derived analytical conditions for cooperation persistence in a system of considerable complexity.

Our IB-model is similar to a previous model considered by Borenstein et al. [30]. However, we were not constrained to a population at carrying capacity, nor assuming simultaneous death and birth events. Instead of running extensive simulations of the resulting IB-model to determine evolutionary outcomes, based on our approach one can solve the condition *κ*_eff_ = 0 to find the parameter that regions separate producer and free-rider fixation.

One limitation of our model is that we consider public goods that are only subject to decay, and not consumption or uptake. This assumption is certainly justified in certain cases, such as enzyme production and pheromone release, since these compounds are not consumed by other organisms. However, our model also serves as good approximation for the case of uptake as long as the density of cells is approximately constant in space.

Our method is unable to capture how spatial correlations (or a relatedness parameter [17]) changes over time. This drawback can be avoided in two different ways. First, it would be possible to model dynamics as an IB-model, make use of an effective mean field correlation theory [29], and then formulate equations that not only model average densities (i.e. eq. (8)), but also pair-correlations between cells. Second, an alternative approach would be to replace the IB-model with an off-lattice model, and then derive equations for the cell-densities and pair-correlations [31]. The latter approach could make it easier to infer interaction strengths directly from time-course imaging data.

The discrepancy between results obtained from the IB-model and our theoretical predictions are mainly due to two factors. Firstly, we have assumed in our analysis of the IB-model that the separation in time scale between PG dynamics and cell division is complete. Although the separation is large (*α/δ ≈* 10^*–*3^), this assumption still introduces some error. Secondly, our analytical results are only correct to first order in 1*/N*. Higher order terms (of order 1*/N* ^3^) could be responsible for the observed discrepancies.

We note that the population dynamics of our IB-model can be predicted with high accuracy with the logistic system (4), which captures the benefit to self experienced by producer cells, but disregards spatial correlations between cell locations. This suggest that cell assortment plays a minor role in determining population dynamics. This is in contrast with other studies [32], and could be due to the lack of mechanical interactions in our model. The above mentioned off-lattice techniques could possibly reconcile these contrasting results. It should also be noted that initial heterogeneity in the cellular distributions (e.g. complete separation between the two cell types) might require more detailed treatment of how spatial correlations evolve in time and space.

Our predictions can be tested experimentally in several ways: (1) either by measuring short-term growth rates as a function of the producer density, or (2) in the long-term, by measuring the fixation probability as a function of the public goods benefit. The latter approach can be modified by changing the background nutrient or serum concentration, as done by Archetti et al. [11]. Another approach would be to track the number and position of single cells and from that calculate effective interaction parameters [33]. This could reveal if, as we claim, spatial correlations play a minor role in determining growth rates, or if interactions and hence average growth rates evolve over time.

In conclusion we here developed a theory of diffusible public goods and producer-free-rider co-evolution, which enabled us to predict the outcome of this complex dynamical process based on the physical parameters of the system. Our results advance understanding of evolutionary dynamics of cooperation in microbial systems and co-evolving cell populations that depend on synergistic signaling, such as observed in biofilms and possibly in tumors.

## Supporting information

Suppl. Methods and Proofs

