## Supplementary material for "Persistence of cooperation in diffusive public goods games": Suppl. Methods and Proofs

### Supplementary Methods

Here we describe the individual-based (IB) model in full detail, and derive the continuum limit of the individual-based (IB) model, which consists of a coupled system of two PDEs and one ODE. We also prove exact results about the growth rates of producers and free-riders, and show how these results apply to a well-mixed version of the IB-model and closely predict the dynamics of the IB-model with limited dispersal. We here present the proofs to all theorems.

Our aim is to derive the growth rates $\Gamma_c$ and $\Gamma_r$ of producers and free-riders respectively, under the assumption of finite cell-size and an uncorrelated population structure. With these growth rates we can predict the evolutionary dynamics of the system using a system of coupled logistic equations. However, in order to obtain the growth rates we first need to consider the continuum limit of the IB-model and show how the density of producer and free-rider cells affect their respective growth rates. We then apply that method to population densities which reflect the discrete nature of the cells in the IB-model, which finally makes it possible to calculate the growth rates in the IB-model with long range dispersal.

#### 1 Individual-based model

We consider a 2-dimensional spatial domain of linear size $L$, discretized into a lattice with $N \times N$ lattice points. The cells are located on this lattice, and each lattice point can be in three states: (i) contain a producer cell, (ii) contain a free-rider cell or (iii) empty. We index the lattice points with integers $i, j = 0, ..., N-1$ and denote the spatial distribution of producer cells with $c_{i,j}$, where $c_{i,j} = 1$ if a producer cell is located at $(i, j)$ and zero otherwise. The free-riders a denoted by $r_{i,j}$. Both cell types have the ability to divide, move and die, processes which will be described in detail below.

The discrete part of the model is coupled to continuous field which describes the public goods concentration, denoted $g(x, t)$. We assume that the public goods is produced by producer cells at some rate $\rho$, diffuses within the domain with diffusion coefficient $D$, and decays at a constant rate $\delta$. This implies that the PG obeys the reaction-diffusion equation

$$\frac{\partial g(x,t)}{\partial t} = D\nabla^2 g(x,t) + \rho c(x,t) - \delta g(x,t) \tag{1}$$

where $c(x,t) = 1$ on lattice points that contain producer cells and $c(x,t) = 0$ otherwise. In order to accurately describe the dynamics within a confined domain, such as a Petri dish, we impose zero-flux boundary conditions on $g(x,t)$, i.e. $\nabla g(x,t) \cdot \mathbf{n} = 0$, where $\mathbf{n}$ is the normal vector of the domain boundary.

The public goods is assumed to affect the rate of cell division of both cell types in an identical fashion and for simplicity we assume a linear relationship between PG concentration and division rate. This implies that the **cell division** rate of a free-rider cell located at position $x$ is given by $\lambda_r = \alpha(1 + g(x,t))$, where $\alpha$ is the baseline rate of cell division in all cells. Producer cells have the same rate of division, but also pay an additional cost $\kappa$, which means that the rate of division for a producer cell located at $x$ is given by $\lambda_c = \alpha(1+g(x,t)) - \kappa$. Upon cell division a neighboring lattice point (using a von Neumann neighborhood) is chosen uniformly at random. If the lattice point is empty the daughter cell is placed there, but if it is occupied cell division fails. **Cell migration** occurs at a rate $\nu$, which is equal for both cell types. When movement has been initiated a neighboring lattice point is chosen uniformly at random. If the lattice point is empty the cell moves to this new position, but if it is occupied movement fails. **Cell death** occurs at a constant rate $\mu$, and leads to the instant removal of the cell from the lattice.

#### 2 Numerical implementation of the individual-based model

We consider a domain of size $L = 1$ cm, discretized into $N = 200$ lattice points, which yields a cell size of $h = 50$ $\mu$m. A simulation of the model is initialized by first prescribing the initial condition for the cells and the public good. Typically we consider a stochastic initial condition of the following form

$$\begin{aligned}
&\Pr(c_{i,j} = 1 \text{ and } r_{i,j} = 0) = p_c \\
&\Pr(c_{i,j} = 0 \text{ and } r_{i,j} = 1) = p_r \\
&\Pr(c_{i,j} = 0 \text{ and } r_{i,j} = 0) = 1 - p_c - p_r \\
&\Pr(c_{i,j} = 1 \text{ and } r_{i,j} = 1) = 0
\end{aligned}$$

where $i,j = 0,...,N-1$ and $p_c$ and $p_r$ are probabilities that satisfy $p_c + p_r < 1$, fix the average initial density of cells. The domain is assumed to be void of PG initially, i.e. $g(x,0) = 0$ for all $x \in [0,1] \times [0,1]$.

Time is discretized with a time step of $\Delta t = 10^2$ seconds, and each time step $n = 1,...,t_{max}$ the following is carried out:

1. The equation for the public good concentration (1) is solved numerically using an ADI-scheme [4] with space step $h$ and time step $\Delta x$, which yields the current concentration $g(ih, jh, n\Delta t)$ at all lattice points $(i,j)$.

2. All the cells are updated in random order according to:

(a) A cell at lattice site $i$ divides with probability $\alpha(1 + g(ih, n\Delta t))\Delta t$ if it is a free-rider or $(\alpha(1 + g(ih, n\Delta t)) - \kappa)\Delta t$ if it is a producer cell.

(b) The cell moves with probability $\nu\Delta t$

(c) The cell dies with probability $\mu\Delta t$

### 3 Continuum limit of the individual-based model

In order to analyse the dynamics of the individual-based model we will derive a continuum description of the model valid in the limit where cell size tends to zero, which for a fixed domain size corresponds to the number of lattice points $N \to \infty$. For simplicity the analysis of the model will be carried out in a 1-dimensional version, but the results obtained also apply to the two-dimensional individual-based-model, with the difference that the diffusion constant in the 2D system is given by $\nu h^2/4$.

We follow a standard technique [3] and let $c_k(t)$, $k = 1, ..., N$, denote the probability of finding a producers at site $k$ at time $t$, and equivalently let $r_k(t)$ represent the occupation probability of free rider cells. The general strategy is to formulate two coupled master equations for the occupation probabilities at site $k$, which will then be approximated by two PDEs, amenable to analysis.

Let us first consider $c_k(t)$. Which are the processes that affect this quantity at a given site?

1. a neighboring cell might divide placing its offspring in the (empty) considered site (with rate $\lambda_c(k\pm1)/2$)
2. an existing cell can die (with rate $\mu$)
3. an existing cell might move away from the considered site (with rate $\nu/2$ in each direction)
4. a neighboring cell might move into the (empty) considered site (with rate $\nu/2$)

Taking all these processes into account and assuming independence between sites we can write

$$\begin{aligned}
c_k(t+\Delta t) - c_k(t) = &+ \frac{\lambda_c(k-1)\Delta t}{2}(1 - c_k(t) - r_k(t))c_{k-1}(t) + \frac{\lambda_c(k+1)\Delta t}{2}(1 - c_k(t) - r_k(t))c_{k+1}(t) \\
&- \mu\Delta t c_k(t) - \frac{\nu\Delta t}{2}c_k(t)((1 - c_{k-1}(t) - r_{k-1}(t)) \\
&+ \frac{\nu\Delta t}{2}(1 - c_k(t) - r_k(t))(c_{k-1}(t) + c_{k+1}(t)).
\end{aligned}$$

The first two terms are due to cell division, the third term arises from cell death and the last two terms are due to migration events. After rearrangement we can write this as:

$$\frac{dc_k}{dt} = \frac{\nu h^2}{2}((1 - r_k)\tilde{\Delta}c_k + c_k\tilde{\Delta}r_k) + \frac{1}{2}(1 - c_k - r_k)\left(\lambda_c(k-1)c_{k-1} + \lambda_c(k+1)c_{k+1}\right) - \mu c_k$$

where

$$\tilde{\Delta}c_k = \frac{c_{k+1} + c_{k-1} - 2c_k}{h^2}$$

is a discrete approximation of the second derivative, where $h$ is the cell size.

A similar calculation for free rider cells shows that $r_k(t)$ evolves according to

$$\frac{dr_k}{dt} = \frac{\nu h^2}{2}((1-c_k)\tilde{\Delta} r_k + r_k \tilde{\Delta} c_k) + \frac{1}{2}(1-c_k-r_k)\left(\lambda_r(k-1)r_{k-1} + \lambda_r(k+1)r_{k+1}\right) - \mu r_k.$$

We now let the cell size $h \to 0$ and assume that the migration rate scales in such a way that $\lim_{h\to 0} \nu h^2/2 = \Gamma > 0$ is the diffusion constant. In this limit $\tilde{\Delta} c_k \to \frac{\partial^2 c}{\partial x^2} + \mathcal{O}\left(h^2\right)$ and in the same way for $r_k$ and we get the following set of coupled PDEs:

$$\begin{aligned}
\frac{\partial c(x,t)}{\partial t} &= \Gamma\left((1-r)\frac{\partial^2 c}{\partial x^2} + \frac{\partial^2 r}{\partial x^2}c\right) + \lambda_c(g)c(1-(c+r)) - \mu c \\
\frac{\partial r(x,t)}{\partial t} &= \Gamma\left((1-c)\frac{\partial^2 r}{\partial x^2} + \frac{\partial^2 c}{\partial x^2}r\right) + \lambda_r(g)r(1-(c+r)) - \mu r \\
\frac{\partial g(x,t)}{\partial t} &= D\frac{\partial^2 g}{\partial x^2} + \rho c - \delta g
\end{aligned} \tag{2}$$

defined on the interval $[0,1]$ with no-flux boundary conditions. The unusual form of diffusion arises because of size exclusion (i.e. a lattice site can only contain a single cell).

### 4 Analysis of the continuum model

The continuum version of the model is considerably simpler to analyze compared to the individual-based-model. However, in order to proceed we require two additional assumptions. First, we assume that the total density of cells is small, i.e. $c(x) + r(x) \ll 1$. This implies that $c(1-(c+r)) \approx c$, and we can therefore drop the logistic growth term in (2) and replace it by a linear growth term. Second, we know that the dynamics of the public good occurs on a time scale which is must faster compared to the dynamics of the cells. This implies that the public good-concentration is in a quasi-steady state, and at every instant determined by the distribution of producer cells. Formally this limit can be achieved by scaling time with the baseline division rate by setting $\tau = \alpha t$, which results in

$$\frac{\alpha}{\delta}\frac{\partial g(x,t)}{\partial \tau} = \frac{D}{\delta}\frac{\partial^2 g}{\partial x^2} + \frac{\rho}{\delta}c - g.$$

In the limit $\alpha/\delta \to 0$ (i.e. when the rate of cell division is infinitely small compared to public good decay) we see that the public good concentration is given by the solution to the ordinary differential equation:

$$Dg'' + \rho c - \delta g = 0 \tag{3}$$

with boundary conditions $g'(0) = g'(L) = 0$. To summarize, our aim is to obtain exact results for the simplified system (which approximates the dynamics of the individual-based-model)

$$\begin{aligned}
\frac{\partial c(x,t)}{\partial t} =& \Gamma\left((1-r)\frac{\partial^2 c}{\partial x^2} + \frac{\partial^2 r}{\partial x^2}c\right) + \lambda_c(g)c - \mu c \\
\frac{\partial r(x,t)}{\partial t} =& \Gamma\left((1-c)\frac{\partial^2 r}{\partial x^2} + \frac{\partial^2 c}{\partial x^2}r\right) + \lambda_r(g)r - \mu r \\
& Dg'' + \rho c - \delta g = 0
\end{aligned} \tag{4}$$

with initial conditions $c(x,0) = c_0(x)$ and $r(x,0) = r_0(x)$, defined for $x \in [0,1]$ and $t \geq 0$ with no-flux boundary conditions.

### 5 Calculation of growth rates

We will investigate the dynamics of the system (4) at short time scales, with the aim of characterizing the impact of instantaneous populations densities on the growth rate of producer and free-rider cells. Experimentally this corresponds to plating the cells at a given spatial density and measuring the change in the total number of producer cells and free rider cells over the course of a short period of time (e.g. one cell cycle). We start by making the following definition.

**Definition.** *By the growth rate of producer and free-rider cells we mean the quantities* $\frac{dN_c(t)}{dt}$ *and* $\frac{dN_r(t)}{dt}$ *where*

$$N_c(t) = \int_0^1 c(x,t)dx$$

*and*

$$N_r(t) = \int_0^1 r(x,t)dx$$

The growth rates at the beginning of a simulation is summarized in the following theorem:

**Theorem 1.** *Assume that the initial densities $c(x,0)$ and $r(x,0)$ are both in $L^2(0,1)$. Then the growth rates of producer cells and free-rider cells in system (4) at time $t = 0$ are given by*

$$\begin{aligned}
\left.\frac{dN_c(t)}{dt}\right|_{t=0} =& (\alpha - \kappa - \mu)c_0 + \frac{\alpha\rho c_0^2}{\delta} + \frac{\alpha\rho}{2}\sum_{n=1}^{\infty}\frac{c_n^2}{\delta + Dq_n^2} \\
\left.\frac{dN_r(t)}{dt}\right|_{t=0} =& (\alpha - \mu)r_0 + \frac{\alpha\rho c_0 r_0}{\delta} + \frac{\alpha\rho}{2}\sum_{n=1}^{\infty}\frac{c_n r_n}{\delta + Dq_n^2}.
\end{aligned}$$

*where $q_n = 2\pi n$, and $c_n$ and $r_n$ are the coefficients of the cosine Fourier series of $c(x,0)$ and $r(x,0)$ respectively.*

*Proof.* We start by noting that

$$\begin{aligned}\frac{dN_c(t)}{dt} =& \frac{d}{dt}\int_0^1 c(x,t)dx = \int_0^1 \frac{\partial c(x,t)}{\partial t}dx \\ =& \int_0^1 \left(\Gamma\left((1-r)\frac{\partial^2 c}{\partial x^2} + c\frac{\partial^2 r}{\partial x^2}\right) + \lambda_c(g)c - \mu c\right)dx \\ =& \Gamma\int_0^1 (1-r)\frac{\partial^2 c}{\partial x^2} + c\frac{\partial^2 r}{\partial x^2}dx + \int_0^1 (\lambda_c(g)c - \mu c)\, dx.\end{aligned}$$

Now

$$\int_0^1 \frac{\partial^2 c}{\partial x^2}dx = \left.\frac{\partial c}{\partial x}\right|_{x=1} - \left.\frac{\partial c}{\partial x}\right|_{x=0} = 0 - 0 = 0$$

because of the no-flux boundary condition. Using integration by parts twice we also have

$$\begin{aligned}&\int_0^1 r\frac{\partial^2 c}{\partial x^2}dx = \left[r\frac{\partial c}{\partial x}\right]_0^1 - \int_0^1 \frac{\partial r}{\partial x}\frac{\partial c}{\partial x}dx \\ &0 - \left(\left[\frac{\partial r}{\partial x}c\right]_0^1 - \int_0^1 c\frac{\partial^2 r}{\partial x^2}dx\right) = \int_0^1 c\frac{\partial^2 r}{\partial x^2}dx\end{aligned}$$

due to the no-flux boundary conditions. This implies that the terms involving second derivates vanish and we get

$$\frac{dN_c(t)}{dt} = \int_0^1 (\lambda_c(g)c - \mu c)\, dx. \tag{5}$$

A similar calculation shows that

$$\frac{dN_r(t)}{dt} = \int_0^1 (\lambda_r(g)r - \mu r)\, dx. \tag{6}$$

Since $c(x,0)$ and $r(x,0)$ are in $L^2(0,1)$ they can be written as Fourier series

$$\begin{aligned}c(x,0) =& c_0 + \sum_{i=1}^{\infty} c_n \cos(2\pi nx) \\ r(x,0) =& r_0 + \sum_{i=1}^{\infty} r_n \cos(2\pi nx).\end{aligned}$$

We note that if $c(x) = c_m \cos(2\pi mx)$, $m \geq 0$, then a solution to the public good-equation (3) is given by:

$$g(x) = \frac{\rho c_m}{\delta + Dq_m^2}\cos(2\pi mx),$$

where $q_m = 2\pi m$. Since the ODE for the public good is linear, a solution for

$$c(x,0) = c_0 + \sum_{i=1}^{\infty} c_n \cos(2\pi n x)$$

is simply given by the linear combination

$$g(x) = \sum_{n=0}^{\infty} \frac{\rho c_n}{\delta + D q_n^2} \cos(2\pi n x).$$

We are now in a position to evaluate the integrals (5) and (6) at time $t = 0$:

$$\begin{aligned}
\left.\frac{dN_c(t)}{dt}\right|_{t=0} &= \int_0^1 \lambda_c(g(x))c(x,0) - \mu c(x,0)dx \\
&= \int_0^1 \left(\alpha\left(1 + \sum_{n=0}^{\infty} \frac{\rho c_n}{\delta + D q_n^2}\cos(2\pi n x)\right) - \kappa\right)\left(c_0 + \sum_{n=1}^{\infty} c_n \cos(2\pi n x)\right) \\
&\quad - \mu\left(c_0 + \sum_{n=1}^{\infty} c_n \cos(2\pi n x)\right) dx \qquad (7)\\
&= (\alpha - \kappa)c_0 + \frac{\alpha\rho c_0^2}{\delta} + \int_0^1 \sum_{n=1}^{\infty}\sum_{m=1}^{\infty} \frac{\alpha\rho c_n^2}{\delta + D q_n^2}\cos(2\pi n x)\cos(2\pi m x)dx - \mu c_0 \\
&= (\alpha - \kappa - \mu)c_0 + \frac{\alpha\rho c_0^2}{\delta} + \frac{\alpha\rho}{2}\sum_{n=1}^{\infty} \frac{c_n^2}{\delta + D q_n^2},
\end{aligned}$$

where we have used the fact that

$$\int_0^1 \cos(2\pi n x)dx = \begin{cases} 1 & \text{if } n = 0 \\ 0 & \text{otherwise} \end{cases}$$

$$\int_0^1 \cos(2\pi n x)\cos(2\pi m x)dx = \begin{cases} 1/2 & \text{if } n = m \\ 0 & \text{otherwise.} \end{cases}$$

A similar calculation for the free-rider cells shows that:

$$\begin{aligned}
\left.\frac{dN_r(t)}{dt}\right|_{t=0} &= \int_0^1 \lambda_r(g(x))r(x,0) - \mu r(x,0)dx \qquad (8)\\
&= (\alpha - \mu)r_0 + \frac{\alpha\rho c_0 r_0}{\delta} + \frac{\alpha\rho}{2}\sum_{n=1}^{\infty} \frac{c_n r_n}{\delta + D q_n^2},
\end{aligned}$$

which concludes the proof. ■

#### 6 Stochastic initial condition

For the individual-based model we consider the following spatially homogeneous initial condition: given probabilities $p_c$ and $p_r$ such that $p_c + p_r \leq 1$, each grid site is occupied by a producer cell with probability $p_c$, occupied by a free-rider cell with probability $p_r$ and is empty with probability $1 - p_c - p_r$. The continuum limit of this initial condition corresponds to constant densities $c(x, 0) = p_c$ and $r(x, 0) = p_r$, which have all Fourier coefficients equal to zero except the first which is simply given by the constant density. In this case theorem 1 tells us that the growth rates should be independent of the diffusion coefficient $D$ and further, that if the two types are seeded with the same density, then the growth rate of free-riders always is greater than that of producers. From simulations we see that both these things are false, which suggests that the continuum approach is a poor approximation.

The main idea to tackle this problem is to represent a random initial condition in terms of a Fourier series with coefficients that are random variables. By calculating the expected value of these coefficients we can apply the same technique as above and calculate the expected growth rates. We start by describing the initial condition of the system as two dependent stochastic processes $C_N(x)$ and $R_N(x)$, where $N$ refers to the number of lattice sites in the system. We partition the domain $[0, 1]$ into $N$ disjoint intervals of length $1/N$, and define $C_N(x)$ and $R_N(x)$ as Bernoulli processes, taking the values $\{0, 1\}$, and being constant on each interval $[i/N, (i + 1)/N]$, where $i = 0, ..., N - 1$. In other words, for all $x \in [0, 1]$ we have:

$$\begin{aligned}\Pr(C_N(x) = 1 \text{ and } R_N(x) = 0) =& p_c \\ \Pr(C_N(x) = 0 \text{ and } R_N(x) = 1) =& p_r \\ \Pr(C_N(x) = 0 \text{ and } R_N(x) = 0) =& 1 - p_c - p_r \\ \Pr(C_N(x) = 1 \text{ and } R_N(x) = 1) =& 0.\end{aligned} \tag{9}$$

For these stochastic processes we have

$$\mathbb{E}[\int_0^1 C_N(x)^2 dx] = \int_0^1 \mathbb{E}[C_N(x)^2] dx = p_c < \infty$$

and similarly for $R_N(x)$. They are thus square integrable and can therefore be represented as Fourier series [2]:

$$\begin{aligned}C_N(x) = C_0 + \sum_{i=1}^{\infty} C_n \cos(2\pi n x) \\ R_N(x) = R_0 + \sum_{i=1}^{\infty} R_n \cos(2\pi n x),\end{aligned}$$

where equality is in the mean-square sense. The coefficients are random variables defined according to

$$C_0 = \int_0^1 C_N(x)dx$$
$$R_0 = \int_0^1 R_N(x)dx$$

and

$$C_n = 2\int_0^1 C_N(x)\cos(2\pi nx)dx$$
$$R_n = 2\int_0^1 R_N(x)\cos(2\pi nx)dx$$

for $n \geq 1$. We are now in a position to state the following theorem:

**Theorem 2.** *Assume that the initial condition to system* (2) *is given by the dependent Bernoulli processes described defined in* (9)*, with parameters* $p_c$ *and* $p_r$ *respectively. Then the expected growth rates of producer cells and free-rider cells are given by:*

$$\mathbb{E}\Big[\left.\frac{dN_c(t)}{dt}\right|_{t=0}\Big] = (\alpha-\mu-\kappa)p_c + \frac{\alpha\rho p_c^2}{\delta} + \frac{\alpha\rho p_c(1-p_c)}{N}\sum_{n=0}^{\infty}\frac{1}{\delta + Dq_n^2} + \mathcal{O}\left(\frac{1}{N^3}\right)$$
$$\mathbb{E}\Big[\left.\frac{dN_r(t)}{dt}\right|_{t=0}\Big] = (\alpha-\mu)p_r - \frac{\alpha\rho p_c p_r}{\delta} - \frac{\alpha\rho p_c p_r}{N}\sum_{n=0}^{\infty}\frac{1}{\delta + Dq_n^2} + \mathcal{O}\left(\frac{1}{N^3}\right).$$

*Proof.* The theorem follows from a direct application of the deterministic result in theorem 1 to a stochastic initial condition. In this case the Fourier coefficients are random variables and the growth rates, which are functions of these coefficients, are therefore also a random variables. Taking the expectation of the growth rates and using the linearity of expectation we get:

$$\mathbb{E}\Big[\left.\frac{dN_c(t)}{dt}\right|_{t=0}\Big] = (\alpha-\mu-\kappa)\mathbb{E}[C_0] + \frac{\alpha\rho\mathbb{E}[C_0^2]}{\delta} + \frac{\alpha\rho}{2}\sum_{n=1}^{\infty}\left(\frac{\mathbb{E}[C_n^2]}{\delta + Dq_n^2}\right) \tag{10}$$
$$\mathbb{E}\Big[\left.\frac{dN_r(t)}{dt}\right|_{t=0}\Big] = (\alpha-\mu)\mathbb{E}[R_0] + \frac{\alpha\rho\mathbb{E}[C_0R_0]}{\delta} + \frac{\alpha\rho}{2}\sum_{n=1}^{\infty}\left(\frac{\mathbb{E}[C_nR_n]}{\delta + Dq_n^2}\right). \tag{11}$$

In order to evaluate these expressions we need to calculate the expected values of the Fourier coefficients for $C_N(x)$ and $R_N(x)$. To simplify this calculation we will consider the following zero mean stochastic processes:

$$\tilde{C}_N(x) = C_N(x) - p_c$$
$$\tilde{R}_N(x) = R_N(x) - p_r,$$

with the following joint probability distribution:

$$\begin{aligned}
&\Pr(\tilde{C}_N(x) = 1 - p_c \text{ and } \tilde{R}_N(x) = -p_r) = p_c \\
&\Pr(\tilde{C}_N(x) = -p_c \text{ and } \tilde{R}_N(x) = 1 - p_r) = p_r \\
&\Pr(\tilde{C}_N(x) = -p_c \text{ and } \tilde{R}_N(x) = -p_r) = 1 - p_c - p_r \\
&\Pr(\tilde{C}_N(x) = 1 - p_c \text{ and } \tilde{R}_N(x) = 1 - p_r) = 0.
\end{aligned} \tag{12}$$

These processes are square integrable and we can write

$$\tilde{C}_N(x) = \tilde{C}_0 + \sum_{i=1}^{\infty} \tilde{C}_n \cos(2\pi n x) = C_N(x) - p_c$$
$$\tilde{R}_N(x) = \tilde{R}_0 + \sum_{i=1}^{\infty} \tilde{R}_n \cos(2\pi n x) = R_N(x) - p_r.$$

This implies that for $n \geq 1$ the processes have the same Fourier coefficients and further that the zero order term in the Fourier expansion of $C_N$ is given by $C_0 = \tilde{C}_0 + p_c$ (and $R_0 = \tilde{R}_0 + p_r$). Let us now calculate the expected value of the Fourier coefficients in (10) starting with the zero order terms:

$$\mathbb{E}[C_0] = \mathbb{E}[\tilde{C}_0 + p_c] = \mathbb{E}\Big[\int_0^1 \tilde{C}_N(x)dx\Big] + p_c = \int_0^1 \mathbb{E}[\tilde{C}_N(x)]dx + p_c = p_c,$$

since $\tilde{C}_N(x)$ has zero mean. Similarly we get

$$\mathbb{E}[R_0] = \mathbb{E}[\tilde{R}_0 + p_r] = \mathbb{E}\Big[\int_0^1 \tilde{R}_N(x)dx\Big] + p_r = \int_0^1 \mathbb{E}[\tilde{R}_N(x)]dx + p_r = p_r.$$

Further we get

$$\mathbb{E}[C_0^2] = \mathbb{E}[(\tilde{C}_0 + p_c)^2] = \mathbb{E}[\tilde{C}_0^2 + p_c^2 + 2\tilde{C}_0 p_c] = \mathbb{E}[\tilde{C}_0^2] + p_c^2.$$

Going back to the definition of $\tilde{C}_0$ we can write

$$\begin{aligned}
\mathbb{E}[\tilde{C}_0{}^2] =& \mathbb{E}\Big[\int_0^1 \tilde{C}_N(x)dx \int_0^1 \tilde{C}_N(y)dy\Big] = \mathbb{E}\Big[\int_0^1 \int_0^1 \tilde{C}_N(x)\tilde{C}_N(y)dxdy\Big] \\
=& \int_0^1 \int_0^1 \mathbb{E}[\tilde{C}_N(x)\tilde{C}_N(y)]dxdy,
\end{aligned} \tag{13}$$

where $\mathbb{E}[\tilde{C}_N(x)\tilde{C}_N(y)]$ is the covariance of $C_N$. Since the random variables of the Bernoulli process $\tilde{C}_N(x)$ are independent we have that if $x$ and $y$ belong to different intervals $\mathbb{E}[\tilde{C}_N(x)\tilde{C}_N(y)] = \mathbb{E}[\tilde{C}_N(x)]\mathbb{E}[\tilde{C}_N(y)] = 0$ since the mean is zero. Now if $x$ and $y$ lie on the same interval $\tilde{C}_N(x) = \tilde{C}_N(y)$ and the covariance

$$\begin{aligned}
\mathbb{E}[\tilde{C}_N(x)\tilde{C}_N(y)] =& \mathbb{E}[\tilde{C}_N(x)^2] = (1-p_c)^2 p_c + (-p_c)^2 p_r + (-p_c)^2(1-p_c-p_r) + (1-p_c)^2 0 \\
=& p_c(1-p_c),
\end{aligned}$$

according to (12). This means that the covariance in (13) is non-zero only if $x$ and $y$ lie on the same interval. For the double integral this means that we only need to integrate over squares of the form $[i/N,(i+1)/N]\times[i/N,(i+1)/N]$, $i=0,...,N-1$, that lie on the diagonal of the domain of integration, and that on these squares we have $\mathbb{E}[\tilde{C}_N(x)\tilde{C}_N(y)]=p_c(1-p_c)$. This implies that

$$\begin{aligned}\mathbb{E}[\tilde{C_0}^2]=&\int_0^1\int_0^1\mathbb{E}[\tilde{C}_N(x)\tilde{C}_N(y)]dxdy=\sum_{i=0}^{N-1}\int_{i/N}^{(i+1)/N}\int_{i/N}^{(i+1)/N}p_c(1-p_c)dxdy\\=&N\frac{p_c(1-p_c)}{N^2}=\frac{p_c(1-p_c)}{N}\end{aligned}\tag{14}$$

since we are integrating across $N$ squares each with area $1/N^2$. This implies that

$$\mathbb{E}[C_0^2]=\mathbb{E}[\tilde{C_0}^2]+p_c^2=\frac{p_c(1-p_c)}{N}+p_c^2.$$

For the mixed zero order term we have

$$\mathbb{E}[C_0R_0]=\mathbb{E}[(\tilde{C}_0+p_c)(\tilde{R}_0+p_r)]=\mathbb{E}[\tilde{C}_0\tilde{R}_0+\tilde{C}_0p_r+\tilde{R}_0p_c+p_cp_r]=\mathbb{E}[\tilde{C}_0\tilde{R}_0]+p_cp_r.$$

Again going back the definition of the Fourier coefficients we can write

$$\mathbb{E}[\tilde{C}_0\tilde{R}_0]=\int_0^1\int_0^1\mathbb{E}[\tilde{C}_N(x)\tilde{R}_N(y)]dxdy.$$

Since $\tilde{C}_N(x)$ and $\tilde{R}_N(y)$ are independent if $x$ and $y$ belong to different intervals we have that $\mathbb{E}[\tilde{C}_N(x)\tilde{R}_N(y)]=0$ (because they are zero mean processes). If $x$ and $y$ belong to the same interval then

$$\begin{aligned}\mathbb{E}[\tilde{C}_N(x)\tilde{R}_N(y)]=&\mathbb{E}[\tilde{C}_N(x)\tilde{R}_N(x)]=(1-p_c)(-p_r)p_c+(-p_c)(1-p_r)p_r\\&+(-p_c)(-p_r)(1-p_c-p_r)+(1-p_c)(1-p_r)0\\=&-p_cp_r\end{aligned}$$

according to (12). This means that

$$\mathbb{E}[C_0R_0]=\mathbb{E}[\tilde{C}_0\tilde{R}_0]+p_cp_r=\frac{-p_cp_r}{N}+p_cp_r,$$

since the integration is identical to that in (14).

We now move on to the higher order coefficients. We have that

$$\mathbb{E}[C_n^2]=4\int_0^1\int_0^1\mathbb{E}[\tilde{C}_N(x)\tilde{C}_N(y)]\cos(2\pi nx)\cos(2\pi ny)dxdy.$$

For the same reason as above this integral can be reduced to diagonal squares where the covariance equals

$p_c(1-p_c)$. This implies that

$$\begin{aligned}\mathbb{E}[C_n^2] =&4\sum_{i=0}^{N-1}\int_{i/N}^{(i+1)/N}\int_{i/N}^{(i+1)/N} p_c(1-p_c)\cos(2\pi nx)\cos(2\pi ny)dxdy\\ =&4p_c(1-p_c)\sum_{i=0}^{N-1}\frac{\cos\left(\frac{(2i+1)n\pi}{N}\right)\sin\left(\frac{n\pi}{N}\right)\left(\sin\left(\frac{2(i+1)n\pi}{N}\right)-\sin\left(\frac{2in\pi}{N}\right)\right)}{2n^2\pi^2}\\ =&\frac{2Np_c(1-p_c)}{n^2\pi^2}\sin^2\left(\frac{n\pi}{N}\right).\end{aligned}$$

We now expand the sine squared in terms of its Maclaurin series:

$$\sin^2\left(\frac{n\pi}{N}\right)=\frac{n^2\pi^2}{N^2}+\mathcal{O}\left(\frac{1}{N^4}\right)$$

and note that this approximation also holds for $n>N$ because of the periodicity of sine. In conclusion we get

$$\mathbb{E}[C_n^2]=\frac{2p_c(1-p_c)}{N}+\mathcal{O}\left(\frac{1}{N^3}\right).$$

A similar calculation shows that

$$\mathbb{E}[C_nR_n]=-\frac{p_cp_r}{N}+\mathcal{O}\left(\frac{1}{N^3}\right)$$

Adding up all the expected values of the Fourier coefficients now gives the following expected growth rates:

$$\begin{aligned}\mathbb{E}\Big[\left.\frac{dN_c(t)}{dt}\right|_{t=0}\Big]=&(\alpha-\kappa-\mu)\mathbb{E}[C_0]+\frac{\alpha\rho\mathbb{E}[C_0^2]}{\delta}+\frac{\alpha\rho}{2}\sum_{n=1}^{\infty}\frac{\mathbb{E}[C_n^2]}{\delta+Dq_n^2}\\ =&(\alpha-\mu-\kappa)p_c+\frac{\alpha\rho p_c^2}{\delta}+\frac{\alpha\rho p_c(1-p_c)}{N}\sum_{n=0}^{\infty}\frac{1}{\delta+Dq_n^2}+\mathcal{O}\left(\frac{1}{N^3}\right).\end{aligned} \tag{15}$$

and

$$\begin{aligned}\mathbb{E}\Big[\left.\frac{dN_r(t)}{dt}\right|_{t=0}\Big]=&(\alpha-\mu)\mathbb{E}[R_0]+\frac{\alpha\rho\mathbb{E}[C_0R_0]}{\delta}+\frac{\alpha\rho}{2}\sum_{n=1}^{\infty}\frac{\mathbb{E}[C_nR_n]}{\delta+Dq_n^2}\\ =&(\alpha-\mu)p_r+\frac{\alpha\rho p_cp_r}{\delta}-\frac{\alpha\rho p_cp_r}{N}\sum_{n=0}^{\infty}\frac{1}{\delta+Dq_n^2}+\mathcal{O}\left(\frac{1}{N^3}\right),\end{aligned} \tag{16}$$

which concludes the proof. ■

Please note that the results can be simplified by using the formula

$$\sum_{n=0}^{\infty}\frac{1}{\delta+Dq_n^2}=\frac{2\sqrt{D}+L\sqrt{\delta}\coth\left(\sqrt{\frac{\delta L^2}{4D}}\right)}{4\delta\sqrt{D}}\approx\frac{1}{2\delta}+\frac{L}{4\sqrt{\delta D}},$$

since the diffusion time scale $\tau_D = L^2/D$ is typically much larger than the time scale of decay $\tau_\delta = 1/\delta$.

### 7 Dynamical system

If the initial distribution of cells is random according to the Bernoulli processes (9) and daughter cells are dispersed uniformly at random on the lattice, then the uncorrelated structure of the initial condition is preserved for all times. This implies that we can use the expected growth rates (15) and (16) to calculate the expected per capita growth rate of producers and free-riders. First let us denote the average density of producers at time $t$ by $c(t)$ and the average density of free-riders by $r(t)$. These are equal to the occupation probabilities $p_c$ and $p_r$, and we can therefore express the per capita growth rates as

$$\begin{aligned}\Gamma_c(c) =&\alpha + \frac{\alpha\rho c}{\delta} - \frac{\alpha\rho c K}{N} + \frac{\alpha\rho K}{N} - \kappa \\ \Gamma_r(c) =&\alpha + \frac{\alpha\rho c}{\delta} - \frac{\alpha\rho c K}{N},\end{aligned} \tag{17}$$

which are obtained by dividing (15) and (16) with $p_c$ and $p_r$ respectively. Here

$$K = \sum_{n=0}^{\infty} \frac{1}{\delta + Dq_n^2} \approx \frac{1}{2\delta} + \frac{1}{4\sqrt{\delta D}}.$$

is the benefit to self induced by limited diffusion of the public good. It is worth noting that the two growth rates only differ by a constant, i.e. $\Gamma_c - \Gamma_r = \frac{\alpha\rho}{N}K - \kappa$, which can be interpreted as a self-benefit (which scales with the cell size $1/N$) minus the cost of production.

Since cell division in the IB-model only is successful if the chosen lattice site is empty the dynamics of the average densities can be described using a system of coupled logistic equations [1]:

$$\begin{aligned}\frac{dc(t)}{dt} =&\Gamma_c(c)c(1-(c+r)) - \mu c \\ \frac{dr(t)}{dt} =&\Gamma_r(c)r(1-(c+r)) - \mu r,\end{aligned} \tag{18}$$

This formulation disregards any spatial correlations in the IB-model and is therefore only exact for the case of uniform dispersal. However, the agreement between the logistic model and the IB-model is good even for local dispersal.

The behavior of the logistic system is summarized in the following theorem.

**Theorem 3.** *The non-negative steady states of system* (18) *with growth functions* (17) *are given by*

1. *The empty state:* $(c_0, r_0) = (0, 0)$*, which is unstable if* $\alpha > \mu$
2. *The free-rider state:* $(c_1, r_1) = (0, \alpha - \mu/\alpha)$*, which is stable if* $\alpha > \mu$ *and* $\frac{\alpha\rho K}{N} < \kappa$
3. *Producer state I:* $(c_2, r_2) = (\frac{a-b+\sqrt{E}}{2a}, 0)$*, which exists if* $E > 0$ *and* $\frac{a-b+\sqrt{E}}{2a} > 0$*, and is stable if* $\frac{\alpha\rho K}{N} > \kappa$

*4. Producer state II:* $(c_3, r_3) = (\frac{a-b-\sqrt{E}}{2a}, 0)$*, which exists if* $E > 0$ *and* $\frac{a-b-\sqrt{E}}{2a} > 0$*, and is unconditionally unstable.*

*where*

$$\begin{aligned} a =& \frac{\alpha\rho}{\delta} - \frac{\alpha\rho}{N}K \\ b =& \alpha - \kappa + \frac{\alpha\rho}{N}K \end{aligned}$$

*and*

$$E = (a+b)^2 - 4a\mu.$$

*Proof.* The result follows from a linear stability of system (18). To find the steady states we solve the system

$$\begin{cases} F(c,r) = 0 \\ G(c,r) = 0 \end{cases}$$

where

$$\begin{aligned} F(c,r) &= \Gamma_c(c)c(1-(c+r)) - \mu c \\ G(c,r) &= \Gamma_r(c)r(1-(c+r)) - \mu r, \end{aligned}$$

which has the solutions $(c_0, r_0) = (0,0)$, $(c_1, r_1) = (0, \alpha - \mu/\alpha)$, $(c_2, r_2) = (\frac{a-b+\sqrt{E}}{2a}, 0)$ and $(c_3, r_3) = (\frac{a-b-\sqrt{E}}{2a}, 0)$.

As for the stability we calculate the eigenvalues of the Jacobian

$$J(c_i, r_i) = \left.\begin{pmatrix} \frac{\partial F(c,r)}{\partial c} & \frac{\partial F(c,r)}{\partial r} \\ \frac{\partial G(c,r)}{\partial c} & \frac{\partial G(c,r)}{\partial r} \end{pmatrix}\right|_{(c,r)=(c_i,r_i)}$$

and determine their signs.

- For $(c_0, r_0)$ we get the eigenvalues $\lambda_1 = b - \mu$ and $\lambda_2 = \alpha - \mu$. If $\alpha > \mu$ we have $\lambda_2 > 0$ and the empty steady state is unstable.
- For $(c_1, r_1)$ we get the eigenvalues $\lambda_1 = \mu - \alpha$ and $\lambda_2 = \mu/\alpha(b - \alpha)$. If $\alpha > \mu$ we have that $\lambda_1 < 0$ and further $\lambda_2 < 0$ iff $b - \alpha = \alpha - \kappa + \frac{\alpha\rho K}{N} < 0$, which is equivalent to the condition $\frac{\alpha\rho K}{N} < \kappa$. Under those conditions $\lambda_1 < 0$ and $\lambda_2 < 0$, which implies that the free-rider dominated steady state is stable.
- For $(c_2, r_2)$ we get the eigenvalues

$$\begin{aligned} \lambda_1 =& -\frac{1}{2a}\left(E + (a-b)\sqrt{E}\right) \\ \lambda_2 =& -\frac{1}{2a}(b-\alpha)\left(a+b-\sqrt{E}\right). \end{aligned}$$

Now $\lambda_1 < 0$ since $E > 0$ and $\frac{E+(a-b)\sqrt{E}}{2a} = \frac{\sqrt{E}(a-b+\sqrt{E})}{2a}$, where $\frac{a-b+\sqrt{E}}{2a}$ was assumed to be positive. For $\lambda_2$ we note that if $a > 0$

$$a + b - \sqrt{E} = a + b - \sqrt{(a+b)^2 - 4a\mu} > a + b - \sqrt{(a+b)^2} = 0$$

and hence

$$-\frac{a + b - \sqrt{E}}{2a} < 0$$

for $a < 0$ we have

$$a + b - \sqrt{E} = a + b - \sqrt{(a+b)^2 - 4a\mu} < a + b - \sqrt{(a+b)^2} = 0$$

and hence

$$-\frac{a + b - \sqrt{E}}{2a} < 0.$$

This implies that the sign of $\lambda_2$ is determined by the factor $b - \alpha$, which is positive if $\frac{\alpha\rho K}{N} < \kappa$, as argued above.

- For $(c_3, r_3)$ we get the eigenvalues

$$\lambda_1 = -\frac{1}{2a}\left(E + (b-a)\sqrt{E}\right)$$
$$\lambda_2 = -\frac{1}{2a}(b-\alpha)\left(a + b + \sqrt{E}\right).$$

We start by noting that

$$\lambda_1 = -\frac{\left(E + (b-a)\sqrt{E}\right)}{2a} = -\frac{\sqrt{E}\left(b - a + \sqrt{E}\right)}{2a} = \frac{\sqrt{E}\left(a - b - \sqrt{E}\right)}{2a} > 0$$

since we assumed that the steady state was positive. This implies that $\lambda_1 > 0$ and the steady state is unstable.

We have now characterized all the non-negative steady states, which concludes the proof. ■
